# Pangenome-Informed Language Models for Synthetic Genome Sequence Generation

**DOI:** 10.1101/2024.09.18.612131

**Authors:** Pengzhi Huang, François Charton, Jan-Niklas M. Schmelzle, Shelby S. Darnell, Pjotr Prins, Erik Garrison, G. Edward Suh

**Affiliations:** Electrical and Computer Engineering, Cornell University, Ithaca, NY, USA; FAIR, Meta, Paris, France; Department of GGI, UTHSC, Memphis, TN, USA; NVIDIA, ECE, Cornell University, Santa Clara, CA / Ithaca, NY, USA

## Abstract

Language Models (LM) have been extensively utilized for learning DNA sequence patterns and generating synthetic sequences. In this paper, we present a novel approach for the generation of synthetic DNA data using pangenomes in combination with LM. We introduce three innovative pangenome-based tokenization schemes that enhance DNA sequence generation. Our experimental results demonstrate the superiority of pangenome-based tokenization over classical methods in generating high-utility synthetic DNA sequences, highlighting significant improvements in training efficiency and sequence quality.

## I. Introduction

Public availability of genome datasets, such as the Human Genome Project (HGP) [1], the 1000 Genomes Project [2], The Cancer Genome Atlas (TCGA) [3], GenBank [4], the International HapMap Project [5], the Human Pangenome Project [6], and the Telomere-to-Telomere project [7], has been instrumental in advancing genomic research. However, large-scale genome sequencing remains costly due to the sophisticated equipment and computational power required [8], [9].

Synthetic data generation offers a scalable alternative for genomic research. Specific tasks such as De Novo genome assembly [10]–[12] and genotype imputation [13] inherently involve the generation of unknown sequences, making them also suitable applications for synthetic data. A good generative model can significantly improve their accuracy and efficiency by predicting missing or incomplete segments.

Deep learning models are widely used in different tasks, even in processing genome sequences and related data [14]– [17]. Although generative adversarial networks (GANs) have been explored for synthetic genome generation, their output is limited to short sequences [18], [19]. LMs have shown their capability to generate synthetic natural languages that are almost indistinguishable from real data. The generated language text can be used to train other models [20]–[22], including those in the medical domain [23], [24]. Proven to be extraordinarily good at processing human language, LMs can also interpret and generate broader text, such as code for programming tasks [25], thereby pushing the boundaries of their application beyond strictly spoken languages.

### The Critical Challenge: DNA vs. NLP Tokenization Differences

While LMs present a promising alternative to understanding and generating long synthetic DNA sequences, effective tokenization of DNA sequences is crucial to leveraging LMs. Applying LMs to DNA sequences faces a fundamental tokenization challenge that differs from Natural Language Processing (NLP). Unlike human language, which has natural word boundaries, semantic units, and grammatical structures, DNA is essentially a string with four letters (nucleotides: A, C, G, T) of billions of characters long and without inherent segmentation. Traditional NLP tokenization methods like Byte Pair Encoding (BPE) rely on frequency-based subword identification. Traditional DNA segmentation methods simply segment sequences into individual nucleotides or length *k* substrings. All these methods face a critical limitation as they lack awareness of the biological structure that can show how genomic variations are organized and segmented.

### Missing Opportunity: Structural Information from Pangenome Graphs

Pangenome graphs encode population-level variation patterns by comparing DNA sequences from a whole population and organizing DNA segments into nodes and connections that capture how genetic diversity manifests among individuals (detailed in §II-B). This graph structure naturally identifies biologically meaningful segmentation boundaries within the DNA sequences, offering a principled alternative to tokenizations that treat DNA as undifferentiated character strings. This opportunity has been largely overlooked in current DNA modeling approaches.

Therefore, we propose three novel pangenome graph-based tokenization schemes for LM-based synthetic data generation that leverage the structural information embedded in pangenome graphs to create biologically-informed segmentation. This work presents the first comparative analysis of classical and pangenome-based tokenization schemes for LMs, specifically GPT-2 and Llama, in learning DNA sequence patterns and generating long synthetic sequences. Our findings reveal that the pangenome graph structure embeds significant information that enhances neural networks’ comprehension of DNA sequences, and can in cases reduce training time and improve scalability. Our contributions are as follows.

- **First**, we introduce three novel pangenome graph-based tokenization schemes that leverage genomic structure to provide biologically-informed segmentation, fundamentally different from frequency-based NLP tokenization approaches that ignore structural relationships.
- **Second**, we demonstrate through comprehensive experiments that our tokenization schemes significantly outperform three classical methods in training efficiency, predictive accuracy, and generation quality for both GPT-2 and Llama architectures, establishing clear computational and performance advantages.
- **Finally**, we establish the first systematic evaluation framework for pangenome-informed tokenization in synthetic DNA generation, providing evidence that structural graph information translates into improved biological utility through sequence alignment quality metrics.

The following paper is structured as follows: Section 2 covers background on synthetic genome generation, Section 3 details tokenization schemes, Section 4 outlines evaluation metrics, Section 5 presents experiments, Section 6 discusses related work, and Section 7 concludes with limitations.

## II. Background

### A. Language Models

Large language models are advanced artificial intelligence systems designed to understand and generate language text based on the data on which they have been trained. These models, such as Mistral [26], Anthropic’s Claude [27], OpenAI’s GPT series [28], [29], Google’s T5 [30], Lamda [31] and Gemini [32], Meta’s OPT [33], BLOOM [34] and LLama [35], [36], etc., utilize massive text data to learn patterns, nuances, and complexities of language. LMs can perform a variety of language-related tasks, including answering questions, translating languages, and even participating in casual conversations. Their ability to process and generate coherent and contextually appropriate responses makes them invaluable tools across multiple fields, from customer service and education to creative writing and technical support.

#### Model Choice with Computational Constraints

Due to computational limitations and the need for systematic tokenization comparison, we focus on accessible baseline models (90M parameter GPT-2 and Llama) trained from scratch rather than larger domain-specific DNA models. We selected these architectures for several key reasons: (1)*Tokenization control*: Tokenization flexibility to isolate the effects of different schemes by training from scratch; (2)*Generative capability*: We need autoregressive generation for synthetic sequence generation, whereas many smaller DNA-specific models (e.g., DNABERT-2 [37], GENA [38]) emphasize classifications; (3)*Feasibility*: Changing vocabularies alters the embedding matrix, requiring full retraining to isolate tokenization effects; DNA-specific models capable of generating long sequences exists [39], but the billion-parameter structure makes it computationally too expensive for our academic research.

### B. DNA Sequences and Pangenome Graphs

#### DNA basics

DNA carries genetic information and has four nucleotides: A, C, G, and T. A *genome* is the complete DNA sequence of an organism, billions long in humans. Individuals of the same species share *>* 99% of this string, but small edits (single-base substitutions, insertions/deletions, structural rearrangements) occur throughout. Unlike natural language, DNA has no whitespace or punctuation; any segmentation is our modeling choice.

#### Pangenome graph

A “pangenome” aggregates the genomic content of many individuals of a species, where “pan” means “all”. A *pangenome graph* [40] encodes multiple genomes as a sequence graph, where shared substrings are collapsed into **nodes**, their breakpoints define node boundaries, **edges** connect adjacent fragments to preserve genome continuity, and **paths** trace individual haplotypes. The graph is constructed through whole-genome multiple alignment, which merges identical subsequences across samples into shared nodes and represents each unique fragment as a distinct node (Figure 3). Details of alignments are given in §IV-B.

#### Why it is useful

The graph structure captures population-level variation while avoiding single-reference bias. Many downstream analyses operate on this structure, caring more about which branch an individual takes rather than on raw nucleotide counts. Importantly, this graph-based representation provides a more principled first-level segmentation that mimics natural language structure: just as words and phrases have meaningful boundaries in text, pangenome nodes capture biologically coherent segments where variation arises, mirroring an underlying “biological grammar”. This natural segmentation provides an ideal foundation for tokenization.

### C. Synthetic Genome Sequence Generation using LMs

We generate synthetic genomes with LMs through five steps (See Figure 1): ➀ **Raw data** (§V-A) are collected; ➁ **Tokenization** (§III) converts sequences into model-ready tokens; ➂ **LM training** fits a generative model with next-token prediction; ➃**Generation** (§V-A) samples synthetic sequences from the trained model; ➄ **Downstream tasks** (§IV) evaluations that assess the quality of the generation and how these sequences can be used.

**Fig. 1:**
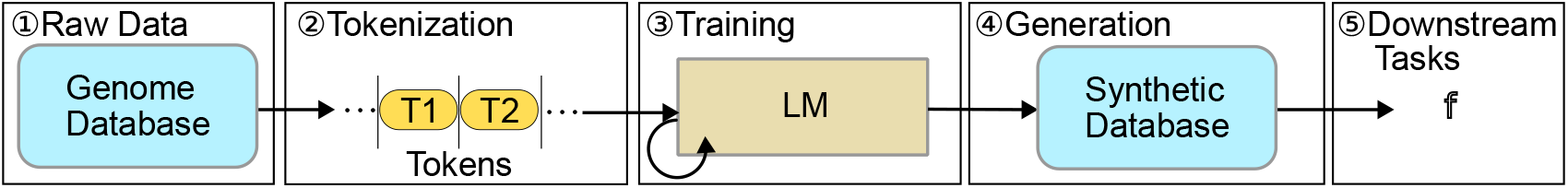
The whole pipeline of synthetic data generation and utilization.

## III. Tokenization of a genome sequence

In this section, we describe the widely used tokenization schemes and our novel schemes. All tokenization schemes are illustrated in Figure 3, where G-headed names denote classical schemes and P-headed names denote graph-based schemes.

### A. Classical tokenizations

#### 1) Genome-based Single Nucleotide Tokenization (GSNT)

GSNT is a straightforward method to tokenize genome sequences applied in prior work([41], [42]). Each nucleotide (A, C, G, T) is treated as an individual token. I.e., “ACGTA” would be tokenized as “A”, “C”, “G”, “T”, and “A”.

#### 2) Genome-based k-mer Tokenization (GKMT)

An alternative is GKMT, where *k*-mers, i.e. substrings of DNA of length *k*, are used as tokens. Depending on the stride, the *k*-mers may overlap or not overlap, and we focus on the non-overlapping *k*-mers in this work. Compared to GSNT, GKMT provides a longer effective context length, but is also highly sensitive to sequence mutations or errors: a single nucleotide insertion or deletion can change all subsequent tokens.

We show how inserting and deletion can affect the GKMT in Figure 2.

**Fig. 2:**
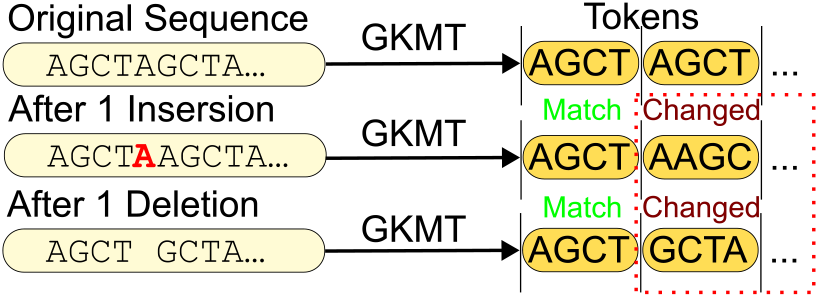
Insertion or Deletion of a sigle nucleotide change all following GKMT (stride equal to *k* = 4) tokens due to the lack of structural awareness of the tokenization scheme.

**Fig. 3:**
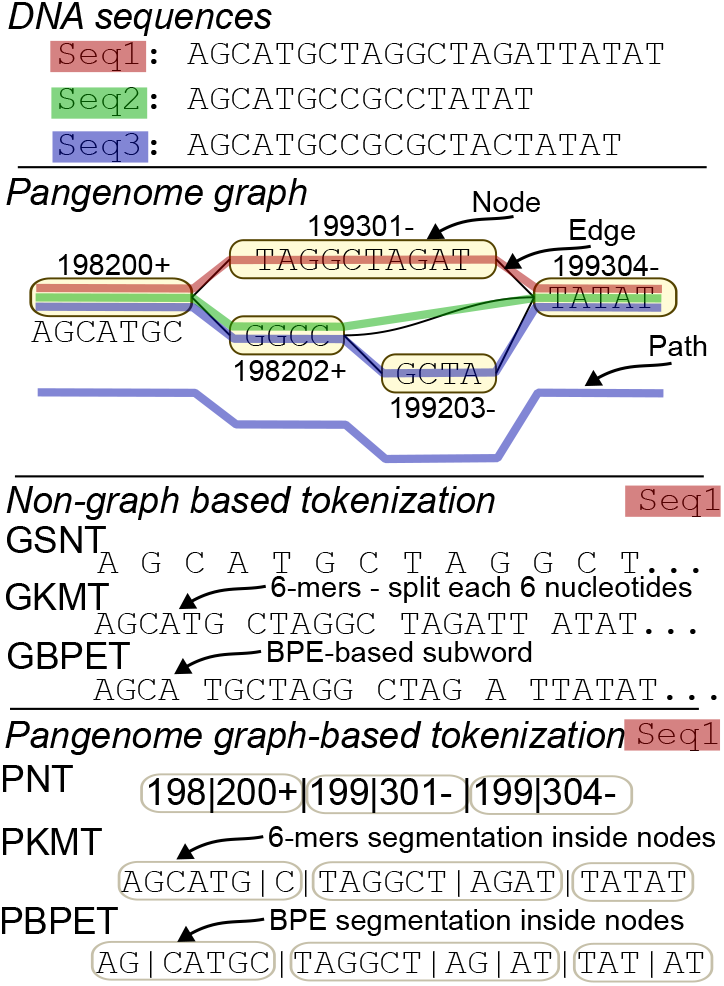
Three DNA sequences and a slice of a pangenome graph with nodes as bubbles (marked with ids), edges as lines. A valid path like the colored routes shows a single individual’s genome. Nodes can be interpreted as forward or reverse orientation, recorded this with +/–. The tokenization schemes output different segmented sequences of the red path, with tokens separated by spaces/bars and nodes by capsule.

#### 3) Genome-based Byte Pair Encoding Tokenization (GBPET)

GBPET, also used in recent studies [37], applies the BPE algorithm [43] to genome sequences. BPE begins with single nucleotide tokens and iteratively merges the most frequent pairs of adjacent tokens to create a vocabulary of longer subword-like tokens. However, BPE training requires too large computational resources if very long DNA sequences are given as inputs. Manual splitting of DNA text is needed in GBPET, which can harm the performance.

### B. Pangenome graph based tokenization

We introduce three novel pangenome graph-based tokenization schemes to overcome these limits:

#### 1) Pangenome-based Node Tokenization (PNT)

The pangenome graph construction assigns each node an abstract numeric ID that encodes its sequence fragment together with its structural context in the graph. Identical sequences occurring at different loci are therefore represented by distinct node IDs. Because nodes encode both the DNA string and its position, identical strings at different locations receive different tokens, producing a very large vocabulary (∼450k in our experiments vs. 50k in common NLP), which presents challenges for model training. We shrink the vocabulary size by splitting the node IDs into two parts (with an additional indicator for reversion, a common variation that causes the sequence to be replaced by its reverse complement). When decoding, tokens “198” and “200+” in Figure 3 are concatenated into “198200+” and decoded as AGCATGC in the forward direction using the pangenome graph’s node-sequence table.

A drawback of PNT is its poor extensibility: adding new sequences requires rebuilding the graph and regenerating IDs, potentially changing the representations largely. We therefore propose the next two schemes not using IDs, and new sequences can be segmented without a complete graph rebuild.

#### 2) Pangenome-based k-mer Tokenization (PKMT)

PKMT first splits sequences at pangenome-graph node boundaries and then cuts each node string into non-overlapping *k*-mers as in GKMT (still *k*=6). Because the graph localizes insertions/deletions to specific nodes, PKMT is more tolerant to variations than GKMT. A main drawback compared with PNT is the loss of explicit positional/structural information from the graph, but using nucleotide strings rather than node IDs provides extensibility.

#### 3) Pangenome-based BPE Tokenization (PBPET)

PBPET first splits sequences at pangenome-graph node boundaries as in PKMT, then runs Byte-Pair Encoding (BPE) that iteratively merges the most frequent adjacent symbol pairs and records the resulting merge rules (the “BPE merges”). We then apply the learned BPE merges to each node sequence, producing variable-length, high-frequency subwords while staying graph-aware via the node pre-segmentation. Unlike GBPET, this initial cut at node boundaries uses and preserves population-level structure from the graph.

## IV. Evaluating synthetic DNA generation quality

A main challenge in proving the utility of our schemes is evaluating the quality of the synthetic genome sequence generation. In our study, we use the prediction accuracy of the model to measure the quality of the generative model. Furthermore, we compare the similarity between synthetic and real genome sequences through sequence alignment.

### A. Model Prediction Accuracy

- **Next token prediction accuracy**: measures how often the model correctly predicts the next token given the correct previous tokens, making it the primary metric for generative models. However, this does not fully reflect sequence accuracy when tokenization is not single nucleotide-based. Predicting “AAAAAC” or “GCTGCT” for the true *k* -mer token “AAAAAA” count both as simply incorrect.
- **Character-level prediction accuracy**: measures the percentage of nucleotides predicted correctly for each token. For example, predicting “AAAAAC” for the true token “AAAAAA” yields an accuracy of 0.83, while predicting “GCTGCT” results in an accuracy of 0. Character-level accuracy measures how well each nucleotide is predicted instead of tokens, making it more consistent and fair across tokenization schemes.

### B. Sequence Alignment Scores

The measurement of similarity between two genome sequences is done using sequence alignment, which is an essential process in many bioinformatic and computational biology tasks. Sequence alignment of DNA involves arranging the DNA sequences to identify regions of similarity. In our case, we use wfmash [44] where the wavefront algorithm [45] is primarily used for pairwise alignment between real and generated DNA sequences. Visualized results (introduced and shown in §V) and multiple scores can be used to evaluate the quality of the alignment.

An example of alignment between a reference sequence and a query sequence is shown in Figure 4.

**Fig. 4:**
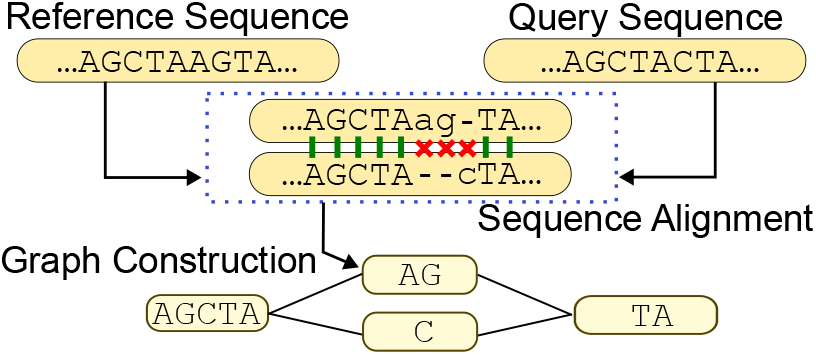
An alignment between two sequence and how it suggests graph nodes. Capitalized nucleotide and green links indicate matches; lowercase nucleotide and red crosses indicate no match; the dashes in the sequences represent the gaps during matching.

An alignment score of 0 indicates no similarity, while a score of 1 represents a perfect match. Take Figure 4 as an example, alignment scores can be defined as:

- BLAST identity (BI): 7*/*10 = 0.7. Defined as the number of matching bases in relation to the number of alignment columns.
- Gap-Compressed Identity (GI): 7*/*9 = 0.78. Counting consecutive gaps as one difference.

The above alignment scores reflect how well the aligned regions match. We also include the **alignment percentage** that measures the proportion of nucleotides in a generated sequence that successfully get aligned to reference genomes. Together, these metrics assess both alignment coverage and quality, providing a measure of biological plausibility and generative fidelity.

#### Why alignment?

We use alignment scores as the primary evaluation metric for synthetic sequences, **rather than short-sequence classification tasks**, for three fundamental reasons:

##### (i) Dataset mismatch for classification tasks

Most datasets of classification tasks, such as those used in DNABERT-2 [37] and GENA [38], evaluate the performance on short sequences (typically 10k or less), focusing on relatively more local characteristics. This length (with possible offset) is too short to build informative pangenome graphs that require long genomic regions of the same species.

##### (ii) Methodological mismatch

Classification tasks emphasize local motifs and short-range patterns, while pangenome graphs capture structural variants and long-range dependencies that are the core advantage of our approach.

##### (iii) Alignment as metric

Alignment scores quantify biological plausibility and consistently track downstream accuracy. Better alignment quality improves SNP/indel calling [46], and increases conserved gene detection, and aids comparative annotation [47]. Alignment-based metrics better match practical genomic goals than generic divergence measures [48]. Prior work in computational biology has used alignment score as a reliable proxy for sequence similarity and utility [49], [50]. In standard pipelines (e.g., reads → reference → variants), higher alignment of generated sequences to real genomes indicates better substitutability for real data (➄ in §II-C).

## V. Experiments

### A. Datasets and LM Choices

In our experiments, we used the human Major Histocompatibility Complex (MHC) region of chromosome 6 as our dataset, extracted from the PanGenome Graph Builder (PGGB) [51] graph of Human Pangenome Reference Consortium (HPRC) year 1 assemblies [6]. Figure 5 shows a simple graph visualization of the data. The MHC region was specifically chosen for its (1) high variation density with complex structural variants essential for pangenome graph construction; (2) long contiguous sequences necessary to capture the structural context that pangenome graphs encode, unlike typical classification datasets using short sequences (∼10 kbp); and (3) population-level diversity across 126 samples with 447 million nucleotides total, providing sufficient variation while maintaining biological realism. The dataset comprises 80% training samples and 20% test samples, with the reference genome temporarily used for hyperparameter tuning before final training.

**Fig. 5:**
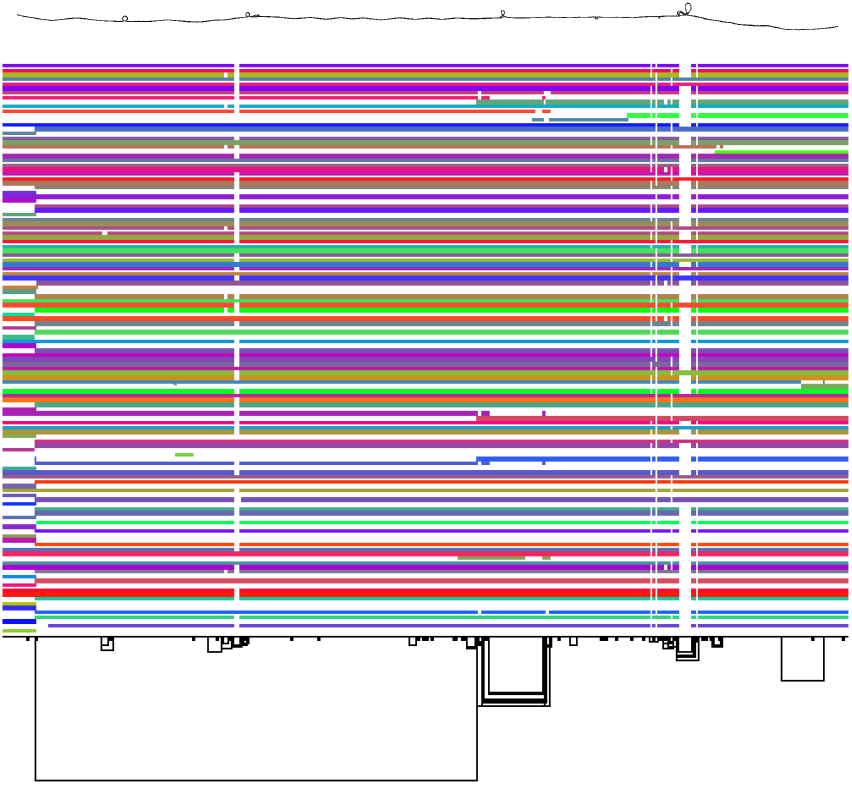
The pangenome graph of the human MHC region of chromosome 6 of the PGGB graph of HPRC year 1 assemblies, with 2D graph visualization (above) and matrix view (below). The circled in the 2D graph and the gaps in the matrix view indicate mutations.

We train all schemes with GPT-2/Llama shown in Table I respectively with Hugging Face 4.24.0 transformers library [52] to isolate tokenization effects. For BPE methods, we set vocabulary size to 4096, following DNABERT2 [37] specifications, and segment long genomes into 10k base pair (bp) sequences due to the computational limits of BPE tokenizer training. To mitigate memorization, we introduce sufficient randomness during generation by sampling with *top-k* = 10, *top-p* = 0.92, and *top-k*_*decend,min*_ = 5, and non-perfect alignment scores (*<*1) indicates that the model is not simply remembering and copying.

**TABLE I:** Configurations used in our experiments.

| Model | Layers | Heads | Hid Dim | FFN | Max Seq |
| --- | --- | --- | --- | --- | --- |
| GPT-2 | 12 | 12 | 768 | 3072 | 1024 |
| LLaMA | 6 | 8 | 768 | 4096 | 2048 |

### B. Experiment Results

We trained the GPT-2 and Llama models on the dataset using four tokenization schemes: GSNT, GKMT, PNT and PKMT. Training was carried out for 90 epochs with a batch size of 16/8 and 1024/2048-token sequences for GPT-2/Llama. The dataset comprises 124 DNA samples totaling 447 million nucleotides. Training time of 90 epochs and vocab size are shown in Table II, obtained on 8 NVIDIA A5500 GPUs. Graph generation costs around 3h on 96-core AMD EPYC 9654. Figure 6 displays token and character-level prediction accuracies. The final accuracies are shown in Table III. PNT not included in the character-level accuracy figures due to the vague definition on predictions and targets with too varied lengths.

**TABLE II:** Vocabulary size and Training time (hours).

| Time | GSNT | GKMT | PKMT | GBPET | PBPET | PNT |
| --- | --- | --- | --- | --- | --- | --- |
| GPT-2 | 56 | 11 | 15 | 17 | 24 | 7 |
| LLaMA | 20 | 5 | 6 | 9 | 12 | 3 |
| Vocab | 4 | 4096 | 5460 | 4096 | 4096 | 2502 |

**TABLE III:** Final accuracy of each tokenization scheme.

| Model | GSNT | GKMT | PKMT | GBPET | PBPET | PNT |
| --- | --- | --- | --- | --- | --- | --- |
| Token Prediction Accuracy |  |  |  |  |  |  |
| GPT-2 | 97.1% | 65.9% | 96.9% | 97.9% | 98.0% | 98.6% |
| LLaMA | 98.7% | 81.8% | 97.7% | 98.5% | 98.6% | 98.8% |
| Character-Level Accuracy |  |  |  |  |  |  |
| GPT-2 | 97.1% | 78.3% | 97.9% | 98.6% | 99.0% | – |
| LLaMA | 98.7% | 85.3% | 98.6% | 99.0% | 99.3% | – |

**Fig. 6:**
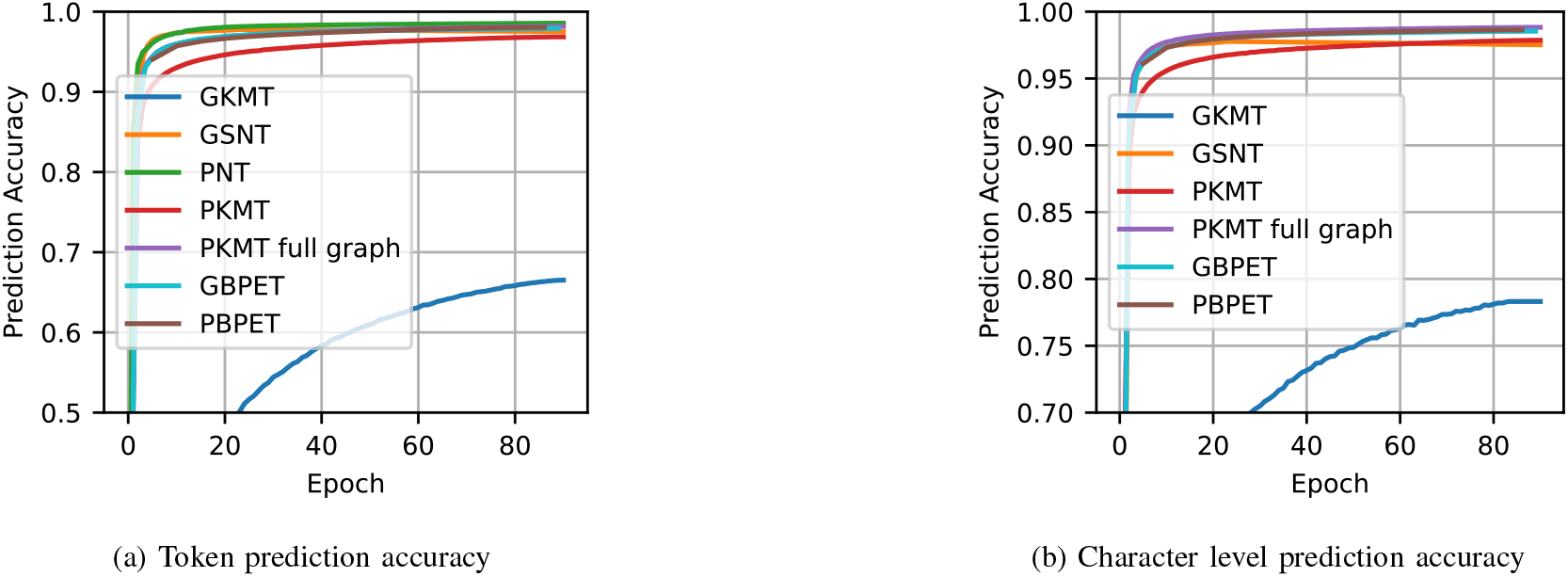
Model prediction accuracies of all tokenization schemes during GPT-2 training.

PNT demonstrated the fastest training time, while GSNT is generally the slowest due to its larger token set. BPE based method is slower than the *k*-mer based method but faster than GSNT. PNT reaches the best peak accuracy the fastest, while GKMT has the worst performance. GSNT initially trains much faster than PKMT for token prediction, but converges to a similar final accuracy. We will see how they perform differently in alignment. Despite having almost the same token tables, we can clearly tell that PKMT’s pangenome graph-aided segmentation helps the model to outperform the one trained by GSNT. The training of the PBPET tokenizer takes around 20 seconds, while the training of GBPET tokenizer takes about 10 minutes, largely due to the larger sequence chunks, and they both have moderate training time.

We report alignment results for GPT–2 generations across all tokenization schemes in Figure 7 (GKMT produces virtually no alignable sequence), and the LLaMA results in Figure 8. The plots show reference coordinates on the *x*-axis and individual generated sequences on the *y*-axis; each dot/segment marks a position in the generated sequence that aligns to the reference. After 90 epochs, only PNT yields long, contiguous alignments for GPT–2. Some sequences show no alignment at all, probably due to stochastic sampling for diversity and occasional misaligned patterns learned during training. LLaMA achieves similar token-level accuracy but exhibits fewer matches overall. With PNT, LLaMA can initiate long runs, yet alignments tend to break off early, especially in high-variation regions (visible as dense dot clusters along the paths). With PKMT or PBPET, LLaMA cannot sustain long aligned sequences like GPT–2.

**Fig. 7:**
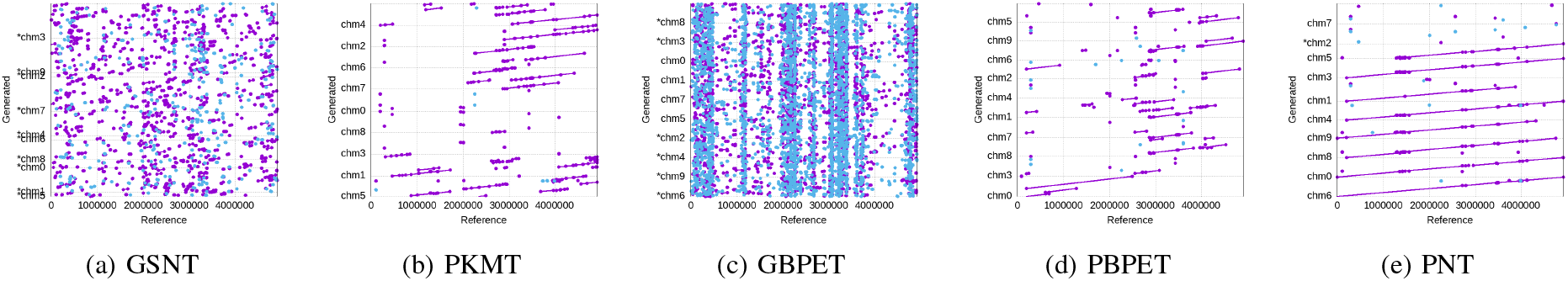
Alignment of a batch of GPT-2 generated sequences against the reference sequence. The X- and Y-axes represent the reference and generated sequences respectively with Longer lines for consistent alignment and dots for short matches.

**Fig. 8:**
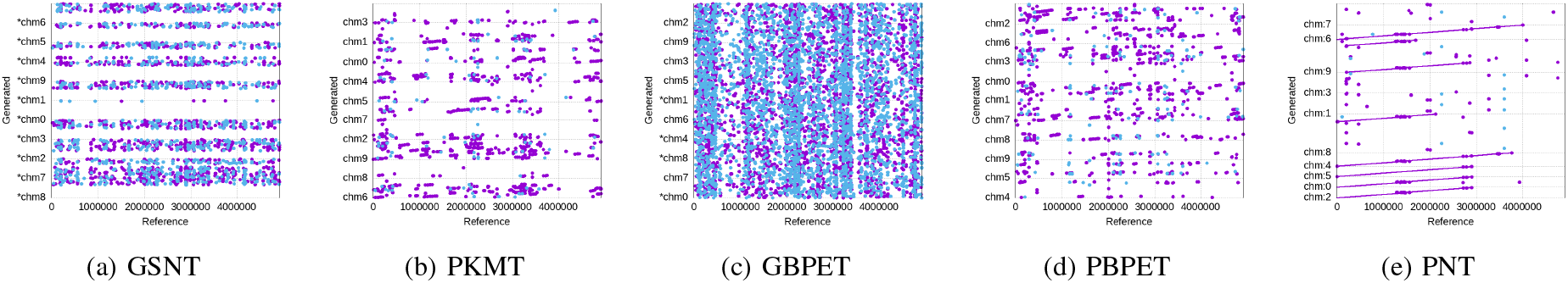
Alignment of a batch of LLaMA-generated sequences against the reference. The X-axis is the reference, and the Y-axis shows the generated sequences. Longer lines indicate consistent alignment, and denser dots indicate frequent short matches.

To quantify generation quality, we show the alignment scores of the generated sequences against the entire dataset (the best match of a query against the entire dataset, GKMT with almost no alignment excluded) in Table IV, with the results for real data as a comparison. A variant-level check is given in §V-C. In addition to GI / BI scores, we show the alignment percentage, indicating the proportion of well-aligned sequences. The segment length refers to the size of the minimizer window during alignment. PNT achieves the highest alignment scores across all segment lengths, while GSNT performs the worst.

**TABLE IV:** Alignment percentages and weighted GI/BI scores of 20 generated sequences per tokenization scheme (∼4M nucleotides each), evaluated across different segment lengths against the original dataset. We report averages over 20 generations, chosen as a practical balance between result stability and computational cost, as larger sample sizes yield similar trends with only marginal variance reduction. Real data metrics are computed using 80% of samples as references and 20% as queries.

| Segment | 1k |  |  | 20k |  |  | 50k |  |  | 200k |  |  |
| --- | --- | --- | --- | --- | --- | --- | --- | --- | --- | --- | --- | --- |
| GPT-2 | Align % | GI | BI | Align % | GI | BI | Align % | GI | BI | Align % | GI | BI |
| GSNT | 81.66 | 0.8712 | 0.9955 | 21.55 | 0.8834 | 0.9893 | 1.42 | 0.8323 | 0.9849 | 0.00 | 0.0000 | 0.0000 |
| PKMT | 52.96 | 0.9443 | 0.9856 | 50.34 | 0.9036 | 0.9932 | 47.87 | 0.8977 | 0.9936 | 8.82 | 0.8656 | 0.9919 |
| GBPET | 71.81 | 0.9873 | 0.9981 | 53.19 | 0.9105 | 0.9931 | 36.93 | 0.9041 | 0.9921 | 0.00 | 0.0000 | 0.0000 |
| PBPET | 44.75 | 0.9081 | 0.9914 | 42.03 | 0.9044 | 0.9943 | 42.29 | 0.9007 | 0.9935 | 9.39 | 0.9029 | 0.9955 |
| PNT | 89.34 | 0.9977 | 0.9999 | 31.01 | 0.9961 | 0.9990 | 33.27 | 0.9920 | 0.9985 | 36.96 | 0.9873 | 0.9982 |
| LLaMA | Align % | GI | BI | Align % | GI | BI | Align % | GI | BI | Align % | GI | BI |
| GSNT | 7.17 | 0.7927 | 0.9906 | 0.00 | 0.0000 | 0.0000 | 0.00 | 0.0000 | 0.0000 | 0.00 | 0.0000 | 0.0000 |
| PKMT | 34.45 | 0.9666 | 0.9960 | 21.19 | 0.8323 | 0.9907 | 6.85 | 0.8232 | 0.9876 | 0.00 | 0.0000 | 0.0000 |
| GBPET | 12.90 | 0.9543 | 0.9870 | 0.00 | 0.0000 | 0.0000 | 0.00 | 0.0000 | 0.0000 | 0.00 | 0.0000 | 0.0000 |
| PBPET | 33.00 | 0.9817 | 0.9969 | 6.41 | 0.8533 | 0.9861 | 5.86 | 0.8356 | 0.9878 | 0.00 | 0.0000 | 0.0000 |
| PNT | 28.80 | 0.9796 | 0.9958 | 5.97 | 0.9970 | 0.9984 | 8.49 | 0.9958 | 0.9987 | 15.02 | 0.9938 | 0.9977 |
| **Real data** | 99.97 | 0.9994 | 0.9999 | 69.23 | 0.9996 | 0.9999 | 61.37 | 0.9991 | 0.9997 | 50.67 | 0.9981 | 0.9993 |

PNT achieves the highest token-level prediction accuracy, while GKMT attains the best character-level accuracy in GPT-2 and performs comparably to PNT in LLaMA. Traditional methods underperform overall, with GKMT reaching below 70% accuracy and GSNT training substantially slower. The performance gap widens in alignment metrics (Table IV), where PNT consistently attains GI and BI scores near 0.99 across segment lengths from 1k to 200k, closely matching real data. Among pangenome-based schemes, PKMT surpasses GSNT in alignment quality and shows potential for long-sequence generation, while PBPET maintains higher stability than its non-graph counterpart GBPET, especially for long segments. In contrast, GBPET performs relatively well only at shorter lengths. Overall, these results highlight the benefits of incorporating pangenome graph structure into tokenization.

LLaMA follows the same general trend as GPT-2 but yields lower generation quality despite higher prediction accuracy and longer prompt length, likely due to the greater degradation with reduced parameter budget dealing with long contexts. We also observe that models can occasionally generate near-empty sequences; this issue can be resolved by adding a brief 20-token prompt during generation starts.

### C. Downstream task: Variant calling

Variant calling evaluates whether the model captures *heterogeneity* (population-level variation) rather than merely memorizing conserved, homogeneous regions. We combined the 20 synthetic sequences with a single training reference path and built a pangenome graph using PGGB [51]. Variants were called for synthetic sequences against the reference path and compared to the “truth” set obtained by calling variants for all other real genomes in the original pangenome graph.

Let *T* be the set of variants from real genomes (ground truth) and *G* the set from generated sequences. After standard normalization (left-normalization, multiallelic splitting, etc.), a *match* is defined when a variant in *G* is equivalent to one in *T* (e.g., same chromosome, position, and REF/ALT, or an accepted equivalence rule).

- **True positives (TP):** variants in *G* matching a variant in *T*.
- **False positives (FP):** variants in *G* with no match in *T*.
- **False negatives (FN):** variants in *T* with no match in *G*.

Precision (Prec), Recall (Rec) and F1 scores are:

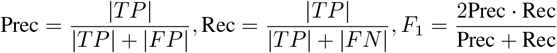

All scores are reported as percentages in Table V. Recall values are modest because only 20 synthetic sequences are contrasted against a truth set derived from 126 real genomes; consequently, many true variants simply never appear in the generated subset. Nonetheless, all pangenome-based tokenizers outperform classical baselines in all metrics, indicating they help the model generate biologically plausible variants (true positives). They more efficiently capture population diversity rather than just conserved sequence. PNT yields lower variant-calling scores than PBPET, reflecting a trade-off between long-range stability and variant richness. PNT produces more coherent and structurally consistent sequences, whereas PBPET captures a greater number of SNPs and indels.

**TABLE V:** Variant-calling performance (precision, recall, and *F*_1_) for each tokenization method.

| Method | Precision (%) | Recall (%) | F1-score (%) |
| --- | --- | --- | --- |
| GSNT | 29.5 | 16.4 | 21.0 |
| PKMT | 65.7 | 28.2 | 39.4 |
| GBPET | 58.1 | 22.9 | 32.8 |
| PBPET | 74.6 | 29.7 | 42.4 |
| PNT | 69.3 | 23.3 | 34.8 |

#### Discussion

To our knowledge, this work is the first to compare the effectiveness of pangenome-based tokenization schemes with classical tokenization schemes for ML learning the pattern of DNA sequences; and also the one of the first to demonstrate the efficacy of LM in generating long DNA.

Our findings reveal that the pangenome graph structure embeds significant and meaningful information, improving neural networks’ understanding of DNA sequences. Our experiments demonstrate how this information can be effectively exploited by graph-based tokenization. The graph-aided segmentation of PKMT/PBPET provides more stable and learnable structural information compared to their classical counterpart, resulting in better overall generation quality. Our results underscore the trade-offs between computational cost and model performance, with pangenome graph-based tokenization schemes showing higher accuracy across tasks. Previous work [6] demonstrates how improved matching is the key point of the pangenome, which “aligns” with our use of the pangenome graph here.

#### Limitations

Our study used smaller models to demonstrate the feasibility of the proposed tokenization scheme. While larger models could yield better results, they also introduce efficiency and resource challenges. Emerging long-context architectures may further enhance performance by capturing long-range DNA dependencies and improving generation consistency. Although pangenome-based tokenization remains advantageous, future work should explore scaling and integrating such architectures to fully assess their potential impact.

## VI. Related work

In this section, we introduce two common genome tasks with the machine learning application.

### A. Classification Tasks

Classification tasks are common in genomics, including:

#### Variant Calling

ML models identify genetic variants such as SNPs and indels in genomes, linking them to diseases or traits. DeepVariant [53], a CNN-based variant caller, out-performs traditional methods, influencing many others [14], [15]. Clairvoyante [54] excels in single-molecule sequencing (SMS), while Clair [55] offers faster RNN-based inference with fewer parameters, without sacrificing accuracy.

#### Gene Expression Analysis

ML models analyze gene expression data to reveal gene-disease relationships. Classical methods like KNN [16], linear/logistic regression [56], and SVMs [57] are used to predict driver genes or cancer risk. CNNs [17], [58] are also applied for cancer classification with RNA-seq data.

Beyond these, CNNs model protein binding [59], cell type identification [60], and non-coding variants [61]. RNNs predict non-coding DNA functions [62] and RNA-protein binding preferences [63]. Transformer models like DNA-BERT [37], [64]–[66] provide strong contextual embeddings for molecular phenotype prediction but face context size limitations due to quadratic scaling. Recent models like Hyena [41] and MambaDNA [42] address these limitations with sub-quadratic scaling for longer contexts. More recent applications of DNA LM like MoDNA [67] for promoter prediction, and GENA [38] for multiple tasks, both use traditional GKMT. Some papers like GPN-MSA [68] for genome-wide variant effect prediction uses GSNT. DNABERT-2 [37] and following work [69] for evolutionary conservation and functional annotation prediction use BPE.

A recent paper [70] presents a similar tokenization approach using pangenome graphs. Although both works independently develop this idea, ours differs by incorporating PNT and PBPET, and focusing on long-sequence generation. In contrast, their work handles shorter sequences (max 5000bp) with node-aided *k*-mer tokenization and focuses on classification tasks.

### B. Generation Tasks

#### Synthetic Data Generation

Synthetic data mimics real data for privacy concerns. GANs have been used for synthetic medical data [18] and DNA sequences coding for proteins [19], though limited by fixed output sizes. Some work [71] utilizes transformers but with limited generation length, and a more recent large model [39] shows generation of submillions in length with a certain level of genomic organization.

#### De Novo Genome Assembly

This involves reconstructing a genome from short DNA fragments without a reference. Deep learning has been applied to de novo peptide sequencing [10]– [12].

## Acknowledgements

This material is based upon work supported by the U.S. National Science Foundation under award No. CCF-2118709 (P.H., J.N.S, P.P., E.G. and E.S.). Any opinions, findings, and conclusions or recommendations expressed in this material are those of the author(s) and do not necessarily reflect the views of the National Science Foundation. The authors also gratefully acknowledge support from National Institutes of Health/NIDA U01DA047638 (E.G.), National Institutes of Health/NIGMS R01GM123489 (P.P. and E.G.).

